# Evaluating Foundation Models for *In-Silico* Perturbation

**DOI:** 10.1101/2025.05.11.653338

**Authors:** Xiong Liu, James Boylan, Theophile Bouiller, Elizaveta Solovyeva, Bulent Ataman, Sebastian Hoersch, Jeremy Jenkins, Murthy Devarakonda

## Abstract

In-silico perturbation (ISP) offers a scalable alternative to traditional gene perturbation experiments, yet evaluation of foundation models for ISP remains underexplored. We introduce a novel evaluation framework, the single cell in-silico perturbation framework (scISP), to benchmark ISP models against in-vitro experimental data using biologically meaningful metrics, including cell state separation accuracy, ISP accuracy, and mean reciprocal rank (MRR) for predicting perturbed genes. Complementary functional analyses evaluate model performance across diverse gene categories. Using scISP, we assess two well-known pre-trained foundation models, Geneformer and scGPT, alongside the deep learning model, GEARS, highlighting their respective strengths and limitations in simulating cell state transitions and identifying perturbed genes. These analyses reveal intrinsic differences across models, offering opportunities to optimize foundation models for specific biological contexts or gene categories. Our extensible framework establishes a robust bridge between computational predictions and experimental validation, advancing gene perturbation research and biological discovery.

## Introduction

Recent advancements in CRISPR-based technologies, such as Perturb-seq^1,2^ and CROP-seq^3^, have revolutionized high-throughput gene perturbation analysis, offering deep insights into gene function and regulatory networks. By combining CRISPR-enabled genetic modifications with single-cell RNA sequencing (scRNA-seq), these techniques enable researchers to study the effects of specific gene alterations on gene expression at the single-cell level. Perturb-seq, in particular, leverages CRISPR interference (CRISPRi) and CRISPR activation (CRISPRa) for targeted gene expression adjustments, thereby enhancing our understanding of gene interactions and regulatory mechanisms.

To complement the capabilities of experimental approaches like Perturb-seq, computational methods have been developed to further analyze and interpret the vast amounts of data generated^4,5^. Foundation models such as Geneformer^6^ and scGPT^7^, based on large language models (LLMs), have shown significant promise in these in-silico experiments. Pre-trained on extensive scRNA-seq datasets, these models accurately represent cellular and gene-related data. They have been successfully applied to various tasks, including correcting batch effects^8^ , classifying cell types^9^, and predicting gene expression profiles^10^, facilitating breakthroughs in understanding cellular processes and disease mechanisms.

Among the various applications of these foundation models, in-silico perturbation (ISP) tasks have emerged as a crucial area of research. ISP leverages computational models to simulate gene perturbations and predict their effects on cellular processes. This approach is not only efficient and cost-effective but also scalable, addressing the limitations of traditional experimental methods. Consequently, ISP has become a focal point for understanding complex biological systems and accelerating the discovery of therapeutic targets and treatments.

Several ISP evaluation methods have been proposed, aiming to predict perturbation responses^11^, identify perturbed target genes^12^, and recover known biological relationships^13^. One particularly promising approach utilizes cosine similarity to measure cell state changes, a method highlighted by Geneformer^6^, which demonstrated significant success in employing foundation models and cosine shift to measure the effects of ISP.

Geneformer utilizes cosine shifts to measure the impact of ISP by identifying genes whose loss causes cells to shift states. For example, it has been used to identify genes whose down regulation shifts cardiomyocytes towards a diseased state^6^. Similarly, scELMO^14^ computes cosine similarity between cell embeddings before and after gene deletion to analyze the targets of in-silico treatment. Another approach, GeneCompass^15^, calculates cosine similarity between post-perturbation and ground-truth cell embeddings, using a mean shift method to measure the effects of ISP. Chen et al.^16^ tested in-silico deletion of genes from the transcriptional regulatory network database (TRRUST) in intestinal fibroblasts from patients with inflammatory bowel disease to determine the cosine shift towards the control intestinal fibroblast state.

Some methods, such as Geneformer^6^ and Chen et al.^16^ applied statistical tests like the Wilcoxon rank sum test to measure cosine shift after gene deletions, emphasizing the importance of rigorous statistical validation in ISP tasks. Although these studies highlight the utility of cosine similarity in evaluating ISP, they often focus on control and disease states and lack comprehensive evaluation against large-scale in-vitro experimental data.

Here, we introduce a novel framework, termed **single-cell ISP (scISP),** for ISP prediction analysis that builds upon the established method of using cosine similarity to measure cell state shifts, while incorporating rigorous statistical tests and a systematic evaluation procedure. While Perturb-seq serves as the primary experimental dataset for this study, the computational framework is broadly applicable to other types of in-vitro perturbation techniques.

The scISP framework consists of the following steps (Figure 1):

**1. Ground Truth Gene Selection:** The process begins with identifying ground-truth genes that exhibit significant changes in expression upon perturbation. This step ensures the biological validity of the data sourced from the Perturb-seq dataset, centered on the K562 cell line^2^ (though similar datasets can be applied). To confirm reliable distinctions between cell states, the E-distance measure^2^ is used to evaluate whether control and target cell transcriptomes differ significantly based on gene expression patterns.
**2. Separation Test (Cell State Validation):** This step validates whether the control and target cell states are statistically distinct and encoded as separable cell representations by the models. For foundation models, these representations exist in an embedding space, while deep learning models like GEARS encode cells based on transcriptomic features. The test evaluates metrics such as within-cluster tightness and between-cluster separation to confirm that the models can differentiate control from target states. Ensuring this distinction is critical for evaluating ISP-generated shifts in step 3, as the cosine shift test measures whether ISP perturbed cells move from the control state toward the target state. If control and target states overlap or are indistinguishable, the concept of shifting between states—and the utility of the cosine shift—is undermined. Thus, this step establishes necessary preconditions for meaningful analysis of the ISP results.
**3. In-Silico Perturbation and Cosine Shift Analysis**: ISP is applied to create computational representations of perturbed cells. The movement of ISP-generated representations towards experimentally validated target cell representations is evaluated by calculating cosine shifts. Statistical methods, including the Wilcoxon rank-sum test and Benjamini-Hochberg correction^17^, are used to assess whether the modeled perturbations significantly shift ISP cells closer to the target cell state. Successful ISP predictions are confirmed when cosine shifts demonstrate consistent movement of the perturbed representation toward the target cell state.
**4. Performance Metrics: Separation Accuracy, ISP Accuracy, and MRR**: To evaluate ISP models, we introduce metrics tailored to distinct aspects of perturbation modeling. Separation Accuracy measures the proportion of ground-truth genes passing the separation test, confirming the model’s ability to encode distinct control and target cell states. ISP Accuracy calculates the percentage of ground-truth genes passing both the separation test and cosine shift analysis, demonstrating successful modeling of perturbation-driven transitions. Mean Reciprocal Rank (MRR) specifically evaluates the quality of predicting the true experimentally perturbed target gene among all input genes passing the separation test, using cosine shift values as ranking criteria. Higher MRR values indicate better prediction performance of the true target gene. Together, these metrics provide a biologically meaningful framework to assess ISP models comprehensively and enable effective model selection.
**5. Functional Analysis and Model Comparison**: Finally, functional analysis examines the biological relevance of genes identified as successfully perturbed across different models. Using Venn diagrams and gene annotations, we analyze the overlap and novelty of functional categories predicted by each model, such as ribosomal genes and translation initiation factors. This comparative analysis highlights each model’s strengths and limitations in capturing biologically meaningful perturbations.

**Figure 1.**
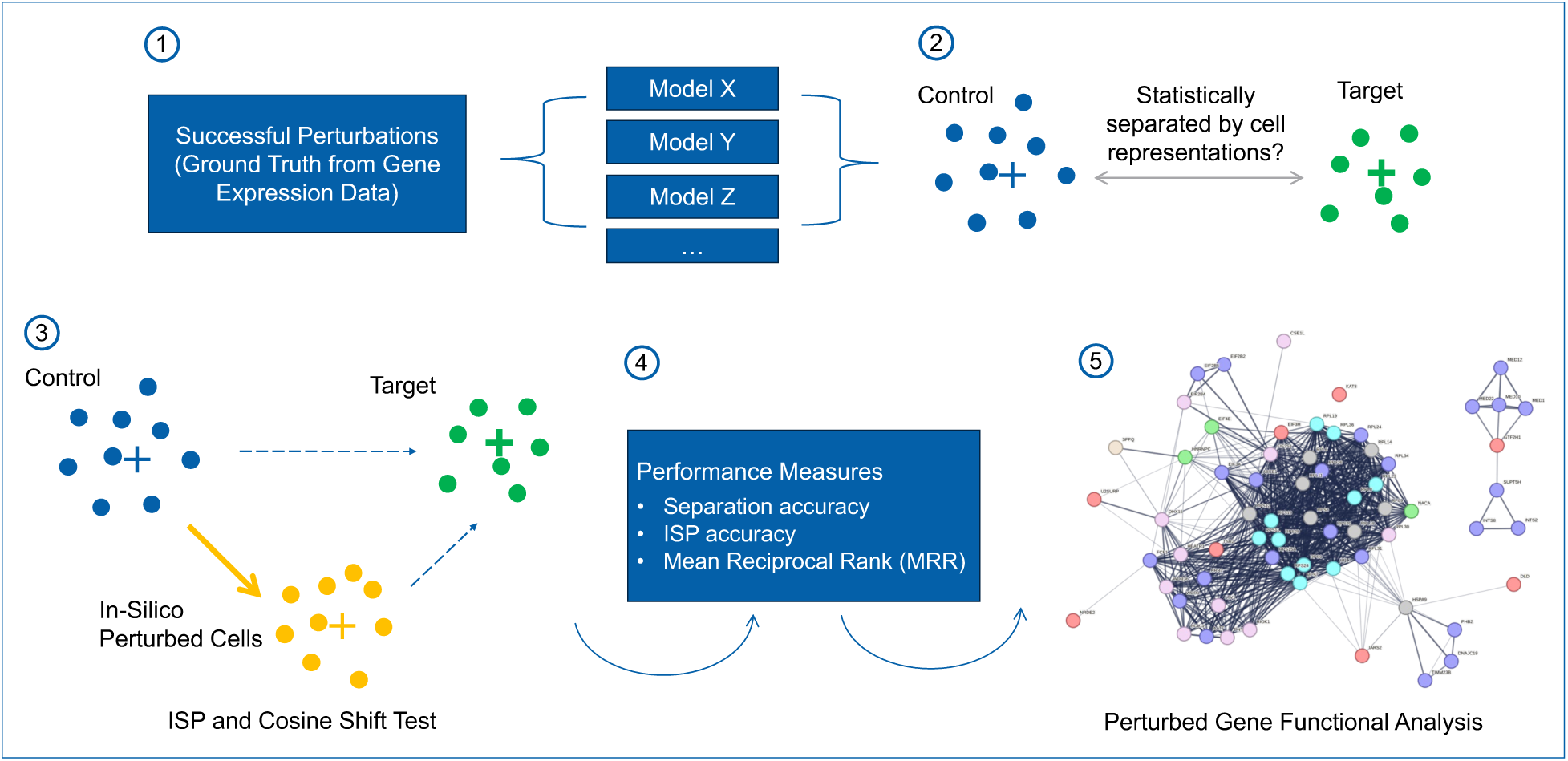
Overview of the scISP Evaluation Framework. The framework consists of five steps designed to evaluate ISP models, including identifying ground-truth genes, validating cell state separation, simulating ISP cell states, assessing model performance with accuracy metrics, and performing functional analysis of perturbed genes.

To demonstrate the utility of the scISP framework, we evaluate foundation models Geneformer and scGPT in their zero-shot versions (i.e., without fine-tuning the embeddings). We also include GEARS^18^, a train-and-test deep learning non-foundation model for ISP that predicts the gene expression changes in perturbed cells. This comparison shows the capability of the framework to handle foundation models (e.g., Geneformer and scGPT) as well as train-and-test models (e.g., GEARS), enabling robust comparison of how well different models perform ISP. In addition, this comparison also highlights the impact of training with the Perturb-seq data.

This work introduces a systematic framework that combines statistical validation of cell state shifts with comprehensive performance metrics. By using large-scale Perturb-seq data and robust statistical tests, this framework addresses gaps in current foundation models benchmarking and ensures *in-silico* perturbations are accurately compared with *in-vitro* experiments, providing an effective methodology for model assessment in the single-cell foundation model field. The framework can produce insights into the ISP capabilities of various models, thus enabling proper choice of models for understanding gene perturbations and cellular responses and supporting therapeutic target discovery.

## Results

### Dataset

To evaluate the performance of foundational models in gene perturbation tasks, we utilized a genome-scale Perturb-seq dataset that employed CRISPR interference (CRISPRi) across millions of human cells^2,19^. Specifically, this study focuses on the Perturb-seq data derived from the chronic myeloid leukemia (CML) K562 cell line. To ensure high-quality data, we applied stringent filtering criteria to remove cells with low numbers of expressed genes, high mitochondrial content, and low unique molecular identifier (UMI) counts. After this quality control step, we retained a total of 310,385 perturbed cells (target cells) with 2,057 unique perturbed genes, as well as 10,305 control cells in the K562 dataset.

To determine successful gene perturbations, we focused on the expression level reductions of the targeted genes under CRISPRi. Using a t-test^20,21^ to compare control and perturbed cells for each targeted gene, success was assessed based on two criteria: 1) A log2 fold change of < −1 in gene expression, and 2) An adjusted p-value ≤ 0.05. Out of the 2,057 perturbed genes in the dataset, 1,740 genes satisfied these thresholds, indicating significant knockdown (Figure 2a).

**Figure 2.**
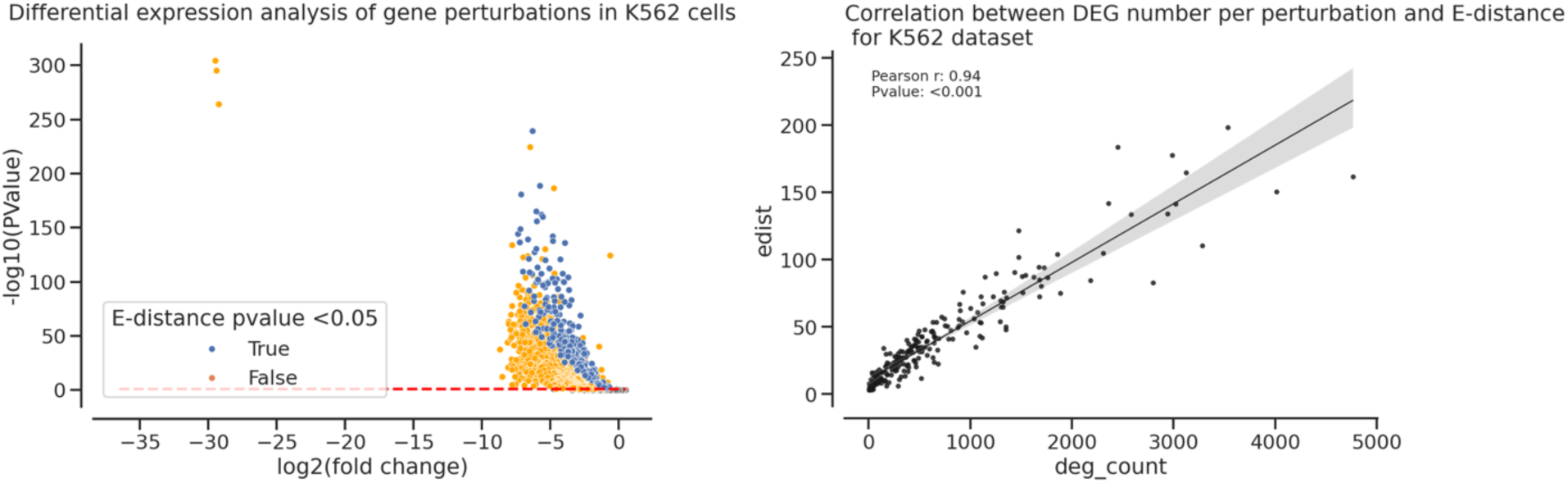
Characteristics of K562 gene perturbations. a (left) Volcano plot illustrating the relationship between log2 fold change and adjusted p-value for perturbed genes in the K562 dataset. Blue markers highlight genes that passed the thresholds for successful perturbations. The dashed line represents the p-value threshold of 0.05. b (right) Scatter plot showing the relationship between the number of differentially expressed genes per perturbed gene and the E-distance score. The strong Pearson correlation (0.94) supports the relevance of E-distance as a metric of transcriptomic impact.

To gauge the broader impact of gene perturbations on the transcriptome, we employed the E-distance metric^2^, a measure of transcriptomic separation between control and perturbed cells. As depicted in Figure 2b, the analysis revealed a highly significant Pearson correlation of 0.94 between E-distance scores and the number of differentially expressed genes. This strong correlation highlights the utility of E-distance as a robust metric for assessing the extent of transcriptomic alterations induced by gene perturbations.

By integrating two criteria—on-target knockdown efficacy (log2 fold change and adjusted p-value) and significant transcriptomic separation (E-distance scores)—we identified 224 perturbed genes as the final set of ground truth perturbations. These genes satisfied both the differential gene expression criteria and exhibited significant transcriptomic differences, as confirmed by their E-distance scores. These 224 ground truth genes represent a curated dataset to evaluate the performance of models.

### Statistical Validation of Cell States

This study evaluates three distinct models within the scISP framework: **Geneformer**, **scGPT**, and **GEARS**, each representing different approaches to in-silico perturbation modeling. Geneformer refers to the pretrained Geneformer 30M 6L model, scGPT corresponds to the pretrained whole-human model, and GEARS represents a comparative deep learning model trained on K562 data using one epoch. These models have distinct methodological approaches, and the flexibility of the scISP framework allows benchmarking across representation strategies.

The separation test, a key step in the scISP framework (Figure 1), validated the ability of each model to distinguish control and target cell states based on gene perturbations. As part of the validation, the test assessed clustering within each model’s cell representation space and determined whether control and target cell states were statistically distinct. Geneformer and scGPT represent cells as embeddings derived from tokenized gene features, while GEARS represents cells using normalized gene expression values, reflecting differences in how models encode cell states within the framework.

The separation test focused exclusively on ISP-applicable genes, meaning validation was performed only on control cells where the experimentally perturbed gene was encoded by the model. Each model applies constraints for how genes are represented in a cell. Geneformer uses up to 2,048 top expressed genes to represent each cell, scGPT includes up to 1,200 non-zero genes with binned features, and GEARS models cells using 5,000 genes. If the perturbed gene was absent from the input lists across all control cells in a dataset, it could not be tested or included in ISP analysis. Such omissions reflect the encoding limitations specific to each model.

Centroid-based cosine similarity combined with one-sided Wilcoxon Rank Sum tests was applied to measure relationships between cell representations. Cosine similarity was chosen because it aligns with the next ISP analysis step involving cosine shifts to assess perturbation effects. For each perturbation gene, control and target cell centroids (average vectors) were calculated in the representation space, and cosine similarity values for individual control cells were measured relative to both the control centroid (“control-to-control”) and the target centroid (“control-to-target”). Similarly, target cells were measured relative to the target centroid (“target-to-target”) and the control centroid (“target-to-control”). The test specifically determined whether control-to-control centroid similarities exceeded control-to-target centroid similarities and whether target-to-target centroid similarities exceeded target-to-control centroid similarities. Genes meeting the threshold of adjusted p-values < 0.05 (Benjamini-Hochberg correction) were considered validated, indicating successful clustering of control and target states. Genes failing validation were excluded from subsequent ISP analyses.

For example, the perturbation gene RPS3 successfully passed validation across all three models, demonstrating clear separation between control and target states. As shown in Figure 3 (Panels a– c), histograms of cosine similarity values confirm that control-to-control similarities were consistently higher than control-to-target similarities, and target-to-target similarities were consistently higher than target-to-control similarities. This reflects distinct clustering patterns for control and target states within each model’s representation space. Additionally, UMAP visualizations in Appendix Figure A further illustrated the cell clustering patterns for RPS3 across the representation spaces of all three models.

**Figure 3.**
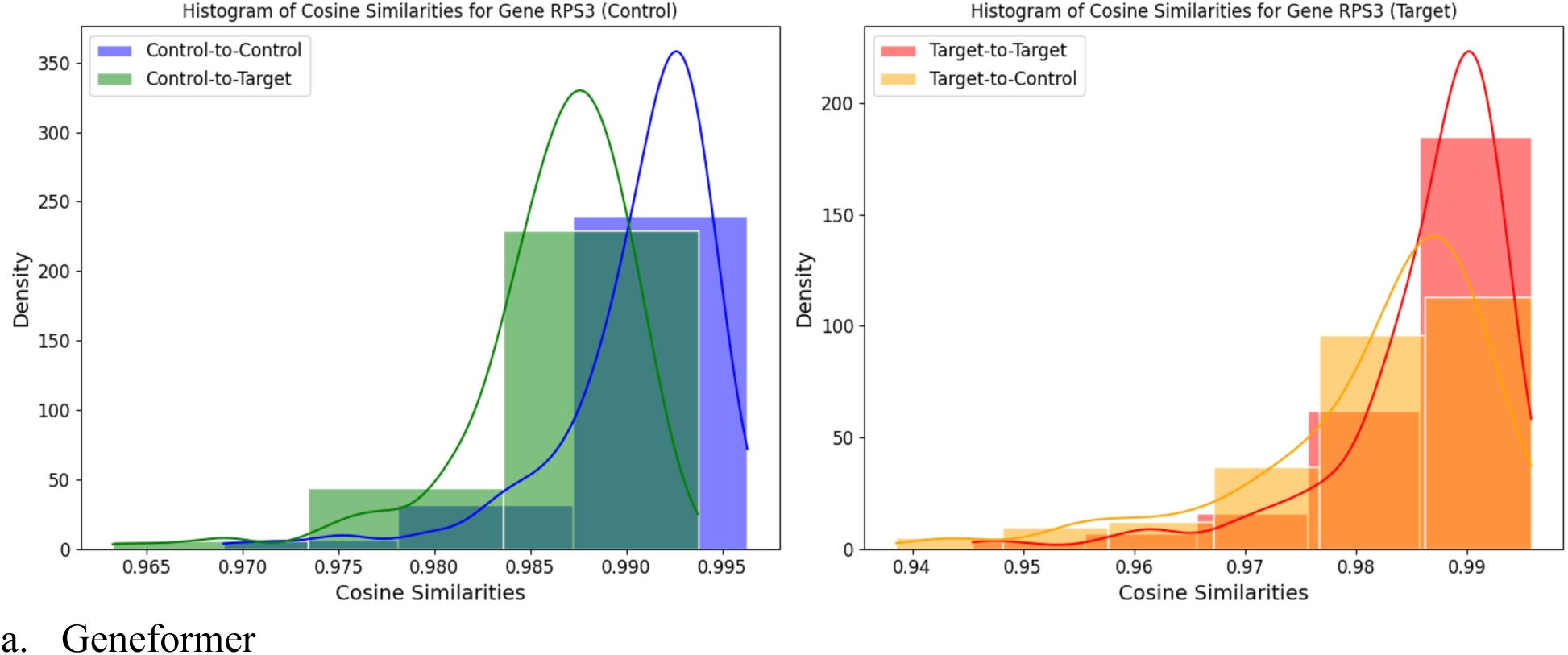
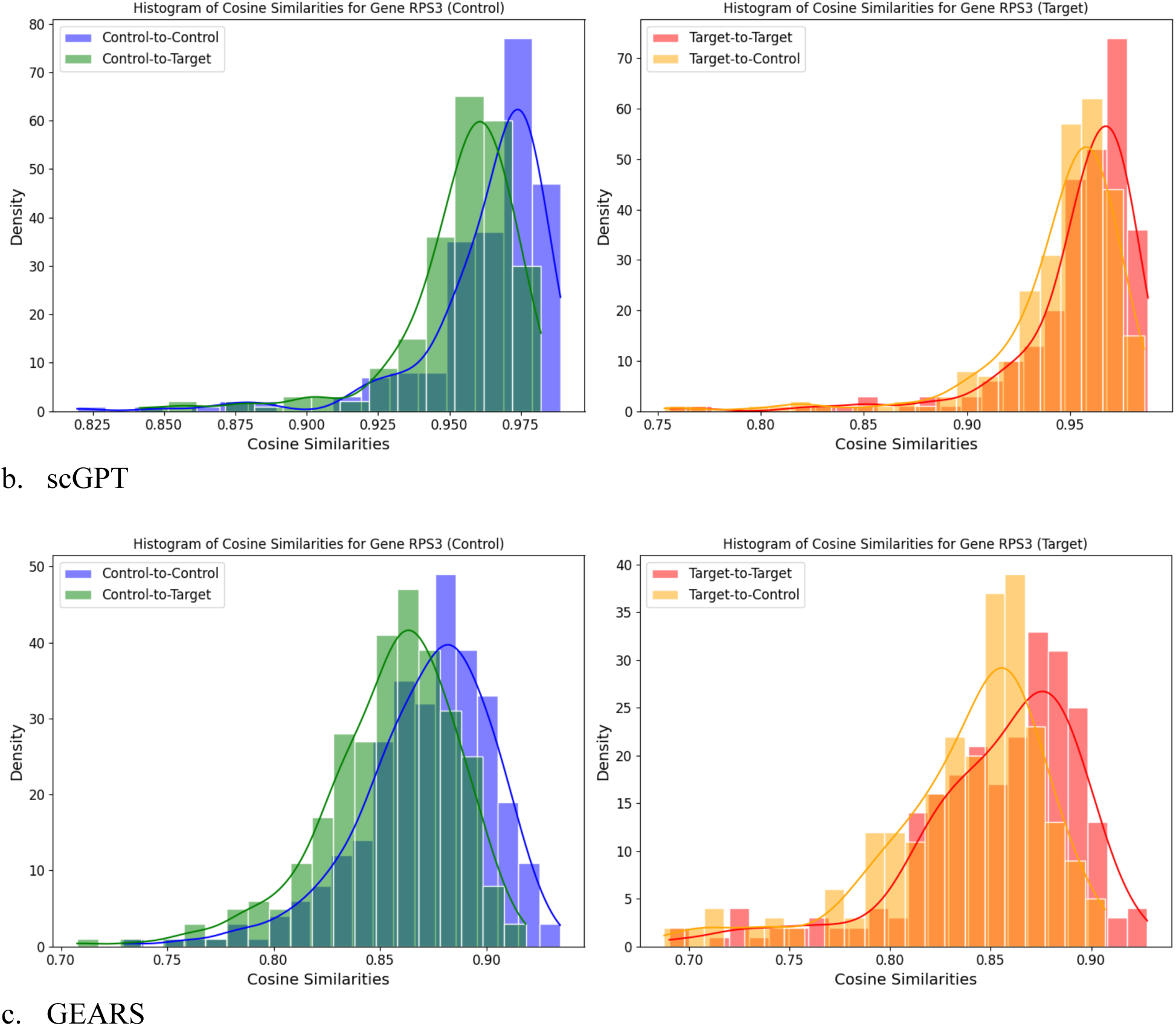
Histograms of Cosine Similarities for Gene RPS3. a) Geneformer, b) scGPT, and c) GEARS models showing control-to-control centroid similarity larger than control-to-target centroid similarity and target-to-target centroid similarity larger than target-to-control centroid similarity, reflecting distinct clustering.

The separation test results for all ground-truth ISP-applicable genes are summarized in Table 1. Among 224 ground-truth genes, GEARS achieved the highest separation accuracy (62.95%), validating 141 genes, followed by scGPT (52.23%), which validated 117 genes, and Geneformer (42.86%), which validated 96 genes. Each model faced limitations due to encoding constraints, which affected the number of ISP-applicable genes. To ensure fairness across models, the separation accuracy was calculated using the total number of ground-truth genes (224) as the denominator. This rigorous validation guarantees that only biologically meaningful perturbations advance to ISP analyses.

**Table 1.** Summary of Genes Passing the Separation Validation Step.

| Model | Ground Truth Genes | ISP Applicable Genes* | Genes Passed | Genes Failed | Separation Accuracy** |
| --- | --- | --- | --- | --- | --- |
| Geneformer | 224 | 217 | 96 | 121 | 42.86% |
| scGPT | 224 | 218 | 117 | 101 | 52.23% |
| GEARS | 224 | 221 | 141 | 80 | 62.95% |

\*ISP Applicable Genes mean the dataset where ISP of control cells can be performed, excluding genes not represented in control cells. \*\*Separation accuracy is calculated as the percentage of ground truth genes that pass the separation validation step.

### In-Silico Perturbation and Quality Measures

The ISP analysis focused on evaluating how effectively each model simulated perturbation effects for the genes that successfully passed the separation test. This step ensures that the models replicate the biological divergence seen in ground-truth perturbations, providing both statistical rigor and comparative benchmarking. For Geneformer, 96 genes passed the separation test, while scGPT validated 117 genes and GEARS validated 141 genes. Only these gene sets were used for ISP evaluation, maintaining methodological consistency across models.

### ISP via Down-Regulation

The log2 fold changes observed in the 224 ground-truth perturbations ranged from −7.36 to −1.08. These values indicate that the experimental perturbations result in gene expression reductions, rather than complete knockouts—roughly 0.0056 to 0.47 times the original expression levels. This down-regulation behavior, more representative of experimental knockdowns than gene deletions, guided the simulation of ISP across all models. Hence, we adopted a down-regulation ISP method, wherein the models simulate a decrease in the perturbed gene’s expression level to align with observed in-vitro perturbations.

### ISP Cell Representation

To generate in-silico perturbed cells, each model employed its own representation strategy. Geneformer was used to simulate down-regulation by moving the perturbed gene/token to the end of the input list for a control cell, rather than removing it entirely as in the original knockout simulations by Theodoris et al^6^. This adjustment reflects the down-regulation behavior observed in in-vitro experimental data. The modified input was then processed through the model to derive ISP cell embeddings. scGPT was used to adjust the binned expression value of the perturbed gene to simulate down-regulation before passing the modified input through the model to obtain embeddings. GEARS was applied to predict normalized expression values for the perturbed cells using graph-based embeddings that capture gene co-expression and perturbation relationships, and the predicted values were used to represent ISP cells.

### Cosine Shift Analysis

The ISP cell embeddings and predicted gene expression values were analyzed by calculating cosine shifts, a metric that quantifies the movement of ISP cells toward the target state relative to control cells. Cosine shifts are calculated as the difference between the cosine similarity of ISP cells to the target centroid and that of control cells to the target centroid. This definition is consistent with Geneformer’s approach of assessing perturbation effects by comparing shifts towards target states. These shifts evaluate the success of ISP in moving perturbed control cells closer to the target cell state.

To assess the significance of cosine shifts, we compared each gene’s ISP cosine shifts (sample A) against a baseline set of random shifts (sample B). Random shifts were generated using control cells and ISP cells from other genes, calculating the cosine shift of each for the gene’s target centroid. This provides a reference distribution of shifts unrelated to the gene’s perturbation. While random shifts are theoretically independent of the ISP, they sometimes exhibited negative medians. To avoid artificial inflation of pass rates for cosine shift tests, baseline shifts were adjusted to use the maximum of the random shifts and zero shifts of the same size as sample A. Statistical significance was determined using the Wilcoxon Rank Sum test, and p-values were adjusted with the Benjamini-Hochberg method to control the false discovery rate (FDR).

### Illustration: RPS3

The perturbation of the gene RPS3 illustrates the methodology and evaluation process. ISP cells for RPS3 were generated from control cells using the respective methods of Geneformer, scGPT, and GEARS. Cosine shifts were calculated for ISP cells relative to control and target cells, and statistical tests validated the significance of these shifts. As shown in Table 2, RPS3 perturbation passed the cosine shift test in all models, demonstrating successful movement of ISP cells closer to the target state. Figures 4A through 4C provide boxplots of cosine shift distributions for RPS3 compared to random shifts, illustrating significant target-centric movement.

**Figure 4.**
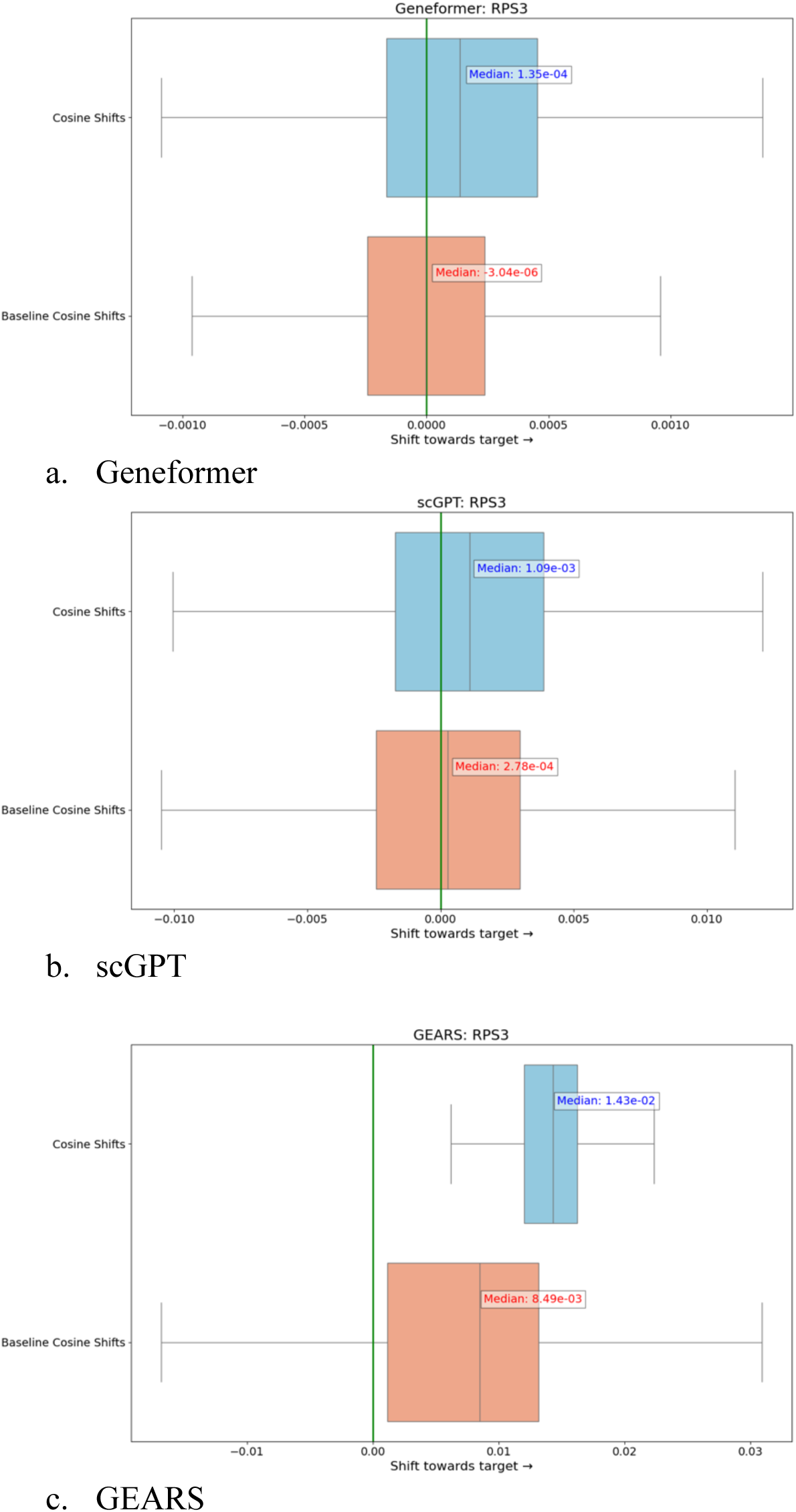
Boxplots illustrating the distribution of cosine shift values for RPS3 and random shifts. a) Geneformer, b) scGPT, and c) GEARS

**Table 2.** Statistical Tests of Cosine Shifts for RPS3 Perturbation.

| Model | Gene | Median Cosine Shift | Median Cosine Shift Random | Rank | p_value adj | ISP Test Success |
| --- | --- | --- | --- | --- | --- | --- |
| Geneformer | RPS3 | 0.000135 | -0.000003 | 2 | 0 | TRUE |
| scGPT | RPS3 | 0.001087 | 0.000278 | 4 | 8.948e-64 | TRUE |
| GEARS | RPS3 | 0.014318 | 0.00849 | 20 | 6.304e-63 | TRUE |

### ISP Accuracy and Predictive Capability

Beyond individual evaluations, Table 3 summarizes the overall ISP accuracy, defined as the percentage of ground-truth genes passing both the separation test and the cosine shift test. GEARS exhibited the highest accuracy at 36.61%, outperforming Geneformer and scGPT, which achieved 14.73% and 14.28% accuracy, respectively. Table 3 also reports MRR (Mean Reciprocal Rank) to evaluate each model’s ability to predict target genes. GEARS achieved better MRR (0.1020) compared to Geneformer (0.0763) and scGPT (0.0638). MRR is a ranking-based metric that measures how effectively the model identifies the true target gene among other genes based on its cosine shift value, with higher MRR indicating better predictive performance. For example, Figure 5 illustrates the ranking of median cosine shifts and the predictive capability of RPS3 among random shifts across Geneformer, scGPT, and GEARS.

**Figure 5.**
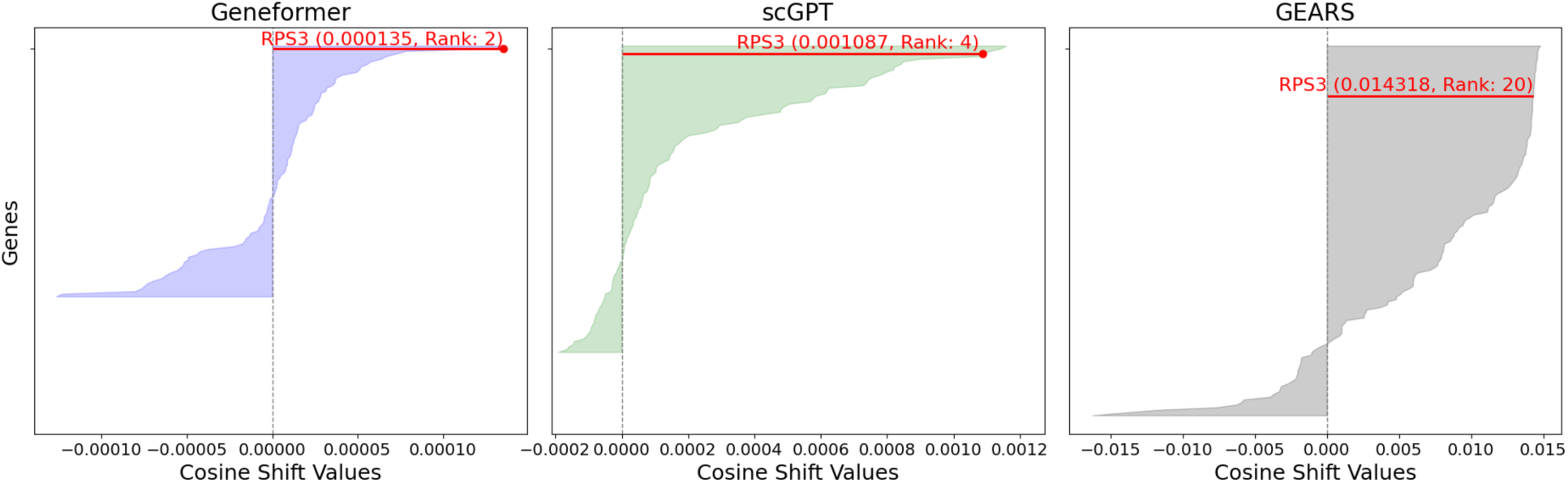
Median cosine shift values and ranks for RPS3 among random shifts across different models.

**Table 3.** Summary of ISP Evaluation.

| Model | Ground Truth Genes | Genes Passing the Separation Test | Genes Passing the Cosine Shift Test | ISP Accuracy* | MRR** |
| --- | --- | --- | --- | --- | --- |
| Geneformer | 224 | 96 | 33 | 14.73% | 0.0763 |
| scGPT | 224 | 117 | 32 | 14.28% | 0.0638 |
| GEARS | 224 | 141 | 82 | 36.61% | 0.1020 |

\*ISP Accuracy denotes the percentage of ground-truth genes passing both the separation validation step and the ISP cosine shift test, demonstrating successful in-silico cell state perturbation. \*\*MRR (Mean Reciprocal Rank) measures the rank of the true perturbed gene among all input genes (restricted to those passing the separation validation step), using cosine shift values as the ranking criterion. Higher MRR values indicate better model performance in identifying the target gene.

This evaluation framework, combining ISP accuracy and MRR, reveals significant differences in model performance. The disparity highlights how representation strategies, perturbation methods, and underlying modeling architectures influence the ability to replicate in-vitro experimental results and prioritize meaningful perturbations. These insights set the stage for further analysis later in the paper.

### Functional Analysis of Correctly Predicted Gene Perturbations

#### Overlap Among Models

To better understand the biological nature of correctly predicted gene perturbations, we analyzed the gene overlap among the three models: Geneformer, scGPT, and GEARS. This evaluation explores how each model identifies perturbed genes, highlighting the differences in their detection capabilities. Figure 6 depicts the overlap in correctly predicted perturbations among the models. Geneformer uniquely identified twelve CRISPRi target genes, scGPT identified seven, and GEARS predicted forty-six genes exclusively. The full list of genes across overlap groups is provided in Appendix C.

**Figure 6.**
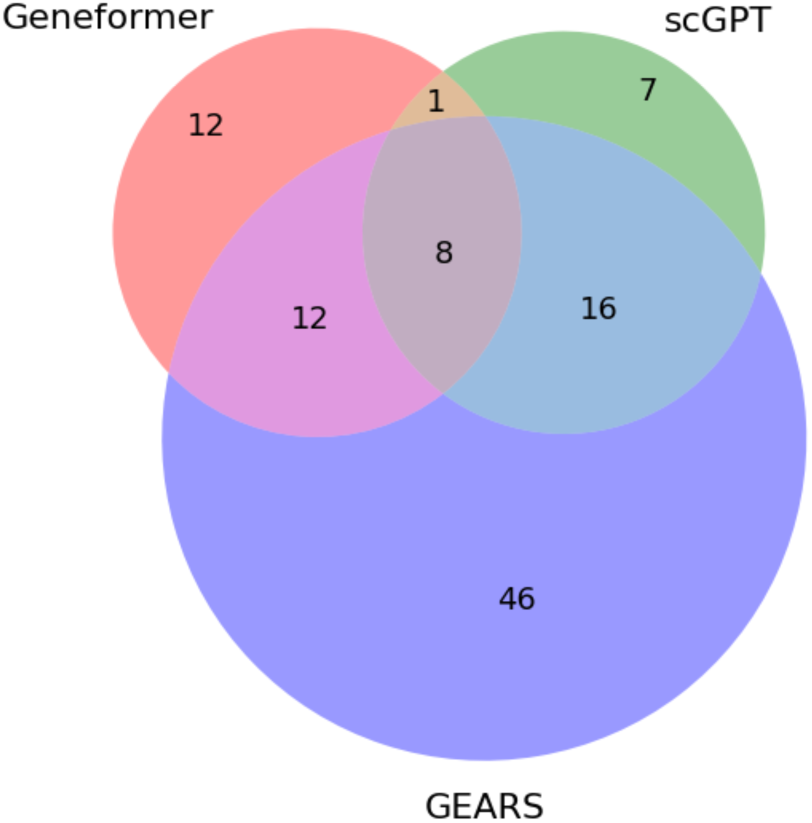
The overlap of correctly predicted perturbations among the models.

#### Functional Categories of Gene Perturbations

To understand the biological roles of genes successfully identified by Geneformer, scGPT, and GEARS, we conducted Gene Ontology (GO)^22^ enrichment analysis using the GO molecular function database and the whole genome as background via the STRING database (string-db)^23^. This analysis revealed significant enrichment in key cellular processes, including ribosomal genes, nucleic acid binding gene sets, translation regulator activities, and protein metabolic processes (the latter derived from GO Biological Processes), as visualized in Figure 7. Statistical details of these results are presented in Supplementary information.

**Figure 7.**
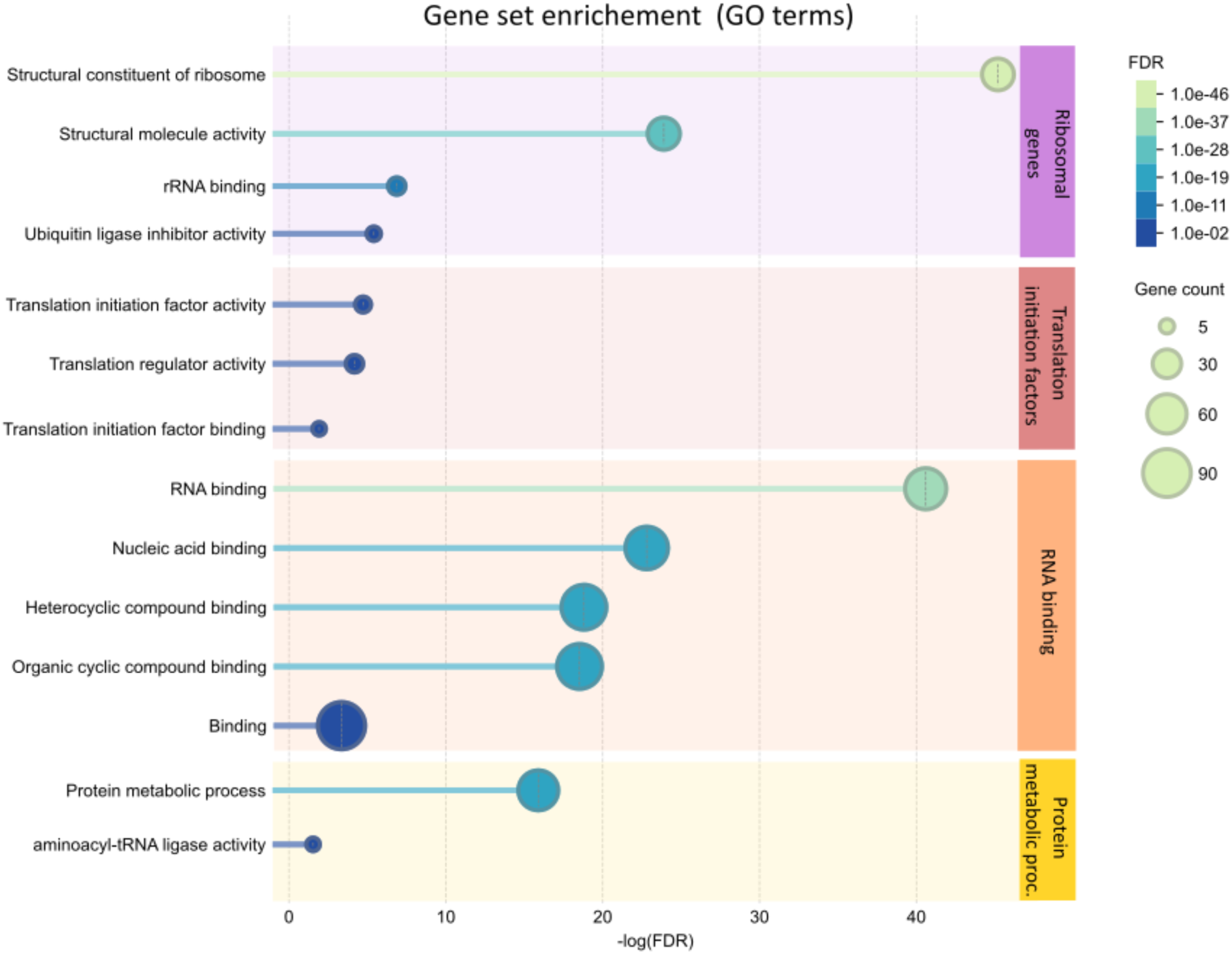
Gene Set Enrichment Analysis for genes identified by at least one model. Enriched categories such as Ribosomal Genes, RNA Binding, Translation Regulator Activities, and Protein Metabolic Processes are grouped based on common molecular functions (GO terms).

These enrichment results highlight fundamental cellular functions of CRISPRi target, emphasizing transcription regulation, protein synthesis, and metabolic pathways. To facilitate downstream analysis, the identified genes were grouped into five broader categories: Ribosomal Genes, Translation Initiation Factors, RNA Binding, Protein Metabolic Processes, and a fifth group for Other Cellular Functions. The grouping was based on general similarity between gene sets (similarity score > 0.7) and overlap of enriched genes (if all enriched genes in one set were fully contained within another, the sets were combined).

Genes assigned to multiple categories were prioritized in a defined order: Ribosomal Genes > Translation Initiation Factors > RNA Binding > Protein Metabolic Processes > Other Cellular Functions, ensuring that genes central to essential processes were categorized appropriately. Table 4 presents the distribution of genes across these functional categories, illustrating trends such as GEARS’ strong propensity for Ribosomal Genes and Translation Initiation Factors, versus Geneformer’s more balanced detection across categories.

**Table 4.**
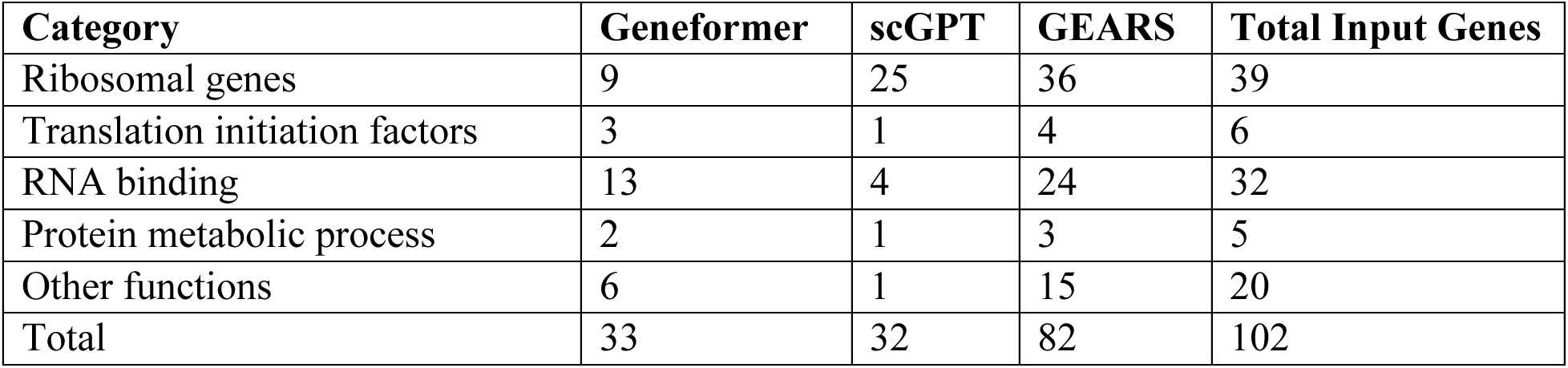
Count of Genes by Category and Model.

| Category | Geneformer | scGPT | GEARS | Total Input Genes |
| --- | --- | --- | --- | --- |
| Ribosomal genes | 9 | 25 | 36 | 39 |
| Translation initiation factors | 3 | 1 | 4 | 6 |
| RNA binding | 13 | 4 | 24 | 32 |
| Protein metabolic process | 2 | 1 | 3 | 5 |
| Other functions | 6 | 1 | 15 | 20 |
| Total | 33 | 32 | 82 | 102 |

#### Gene Interaction Analysis

Interaction networks for correctly identified gene perturbations were explored using string-db. Figure 8 presents a gene-gene interaction map, where nodes represent genes perturbed by at least one model, and edges represent interaction strength among them (based on evidence aggregated from string-db). Node colors correspond to overlap groups (Figure 6): red for Geneformer-alone genes, blue for GEARS-alone genes, and green for scGPT-alone genes. Intermediate colors show shared predictions (e.g., pink for Geneformer and GEARS). Grey nodes represent perturbations shared across all three models.

**Figure 8.**
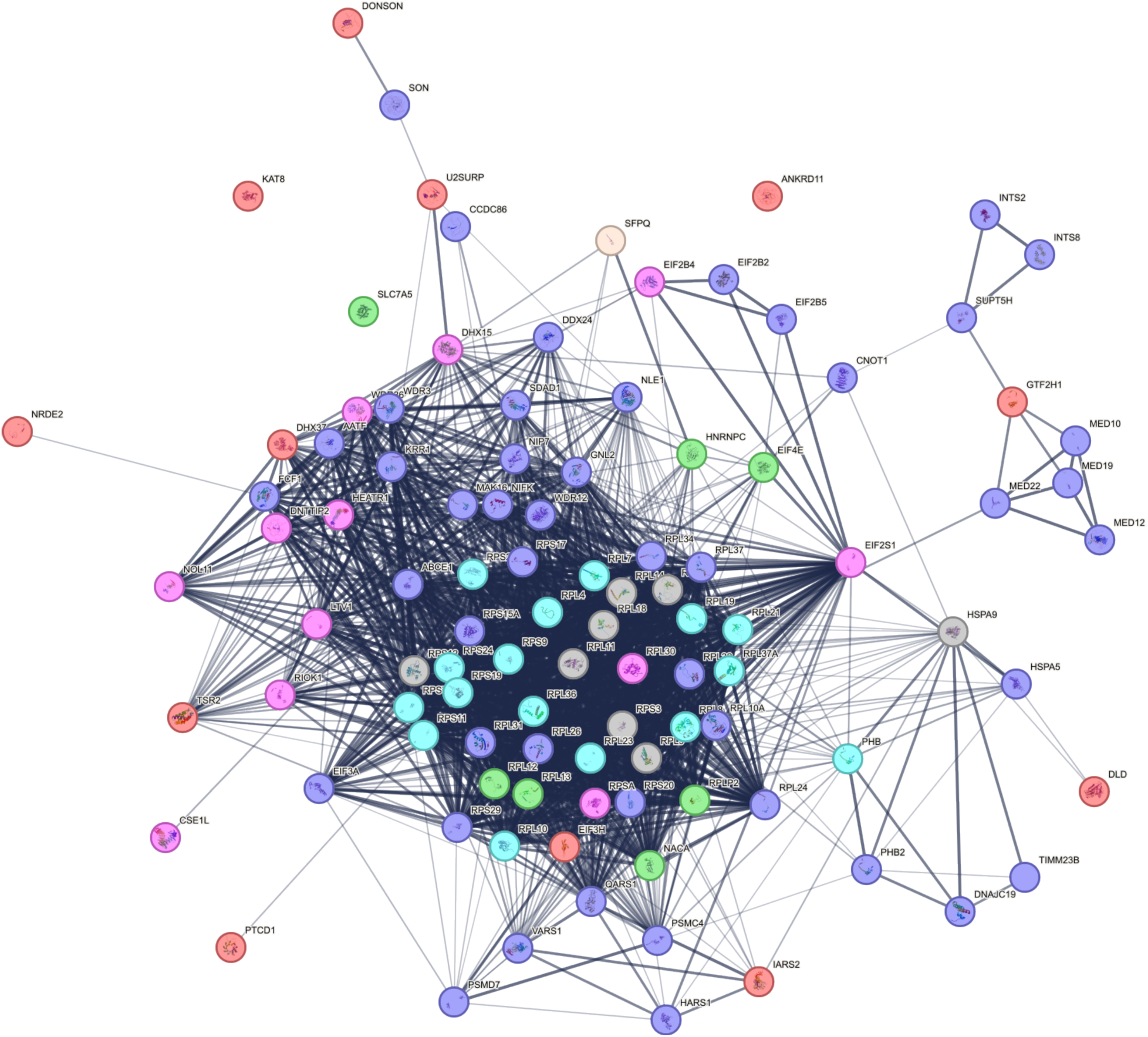
Gene-Gene Interaction Map for predicted genes using string-db evidence. Node colors represent overlaps from Figure 6, and edge thickness corresponds to evidence strength.

GEARS-exclusive genes tended to cluster strongly within enriched categories such as translation initiation and protein metabolic processes, forming distinct and dense interaction networks. By contrast, Geneformer-specific genes showed fewer associations, indicating broader biological detection but less evidence clustering among known cellular processes. To quantitatively assess enrichment across models, we performed further GO enrichment analysis using the entire set of 224 ground-truth genes as the background. Results showed that genes identified by GEARS and scGPT were significantly enriched in structural constituents of ribosomes (GO:0003735), with FDR values of 1.3 × e-3 and 1.7 × e-9, respectively. Conversely, Geneformer did not demonstrate statistically significant enrichment within this category, further distinguishing detection strategies among the models.

#### Predictive Power Across Functional Categories

To evaluate model performance across functional categories, precision and recall metrics were computed as summarized in Table 5. Precision measures the proportion of correctly predicted genes within a category among all successful perturbations, while recall assesses the proportion of correctly predicted genes relative to all ground-truth perturbations within that category.

**Table 5.** Precision and Recall by Model and Category.

| Category | Geneformer Precision | Geneformer Recall | scGPT Precision | scGPT Recall | GEARS Precision | GEARS Recall |
| --- | --- | --- | --- | --- | --- | --- |
| Ribosomal Genes | 0.27 | 0.23 | 0.78 | 0.64 | 0.44 | 0.92 |
| Translation Initiation Factors | 0.09 | 0.50 | 0.03 | 0.17 | 0.05 | 0.67 |
| RNA Binding | 0.39 | 0.41 | 0.13 | 0.13 | 0.29 | 0.75 |
| Protein Metabolic Process | 0.06 | 0.4 | 0.03 | 0.2 | 0.04 | 0.6 |
| Other Cellular Functions | 0.18 | 0.30 | 0.03 | 0.05 | 0.18 | 0.75 |
| **Average (SD)** | 0.20 (0.12) | 0.37(0.09) | 0.20 (0.29) | 0.24 (0.21) | 0.20(0.15) | 0.74(0.11) |

GEARS outperformed other models with higher recall values, particularly for Ribosomal Genes (0.92) and RNA Binding (0.75), indicating its robust ability to detect broader perturbations within critical pathways. Geneformer showed balanced detection across all categories, while scGPT exhibited high precision and recall in Ribosomal Genes (precision 0.78, recall 0.64), highlighting its bias toward detecting perturbations in structural components. This analysis underscores each model’s effectiveness in identifying biologically meaningful perturbations within specific functional categories.

#### Cosine Shift Patterns Across Functional Categories

Boxplots of median cosine shift distributions reveal distinct patterns across functional categories: Ribosomal Genes, Translation Initiation Factors, RNA Binding, and Protein Metabolic Processes (Appendix Figure D). Ribosomal Genes exhibited larger median cosine shifts across all models, demonstrating their distinction from other categories. Conversely, RNA Binding genes, translation initiation factors, and protein metabolic process genes displayed smaller median cosine shifts, suggesting less distinct perturbation predictions relative to ribosomal genes.

## Discussion

In this study, both separation accuracy and ISP accuracy were calculated using the total number of ground-truth genes (224) as the denominator to ensure fairness and consistent benchmarking across models. This approach penalizes models that fail to represent certain ground-truth genes due to input constraints or encoding limitations. By using the full ground-truth dataset, our evaluation avoids inflated scores for models with narrower gene representation while emphasizing perturbation breadth and prediction precision. Models like GEARS, which successfully encoded 221 of the 224 ground-truth genes, applied separation testing and ISP simulation to a larger pool of genes compared to Geneformer and scGPT. This design ensures that performance metrics reflect the sequential filtering process models undergo: ground-truth genes → ISP-applicable representations → validated perturbations → successful ISP predictions. Importantly, ISP simulations are performed only on genes that pass the separation validation step, ensuring they are biologically meaningful and exhibit distinct control-target clustering.

While the main contribution of this work is the systematic benchmarking methodology, the results of applying this methodology to the zero-shot (pre-trained only) Geneformer and scGPT, and to the train-and-test GEARS model warrant discussion. Cell state validation results (Table 1) show that GEARS achieved the highest separation accuracy (62.95%), followed by scGPT (52.23%) and Geneformer (42.86%). ISP results similarly demonstrated that GEARS reproduced more in-vitro perturbations (with ISP accuracy of 36.61% and MRR of 0.1020) than Geneformer and scGPT.

The performance difference is likely due to GEARS’ supervised learning approach, which minimizes the difference between in-silico perturbed cells and target cells, leading to higher cosine similarity and significant cosine shifts. Since we used pre-trained only Geneformer and scGPT, their embeddings do not have the benefit of being trained to distinguish control and target cells in the Perturb-seq dataset. In the future, we propose to fine-tune these models on the Perturb-seq dataset and repeat the analysis.

Further analysis of the cosine shift scores revealed that GEARS generally produced the largest shifts across genes, reflecting its ability to amplify differences between control, ISP, and target states. These cosine-shift scores represent a composite effect: larger **cos(ISP cell, target centroid)** values due to GEARS minimizing the distance between perturbed cells and target states during training, combined with smaller **cos(control cell, target centroid)** values due to GEARS distinguishing control and target cell states more strongly in the separation test. For example, for the gene RPS3, GEARS yielded a median cosine shift of 1.4318e-2, compared to Geneformer’s 1.3548e-4 and scGPT’s 1.0872e-3. Appendix B highlights similar trends across other top-ranked genes by median cosine shift values.

When comparing these cosine-shift magnitudes to previously published results, our values align well with findings in the literature. For instance, the scELMO^10^ study defines a cosine shift score larger than 1e-4 as indicative of a gene’s potential as a therapeutic target. Similarly, Theodoris et al.^6^ reported significant embedding shifts towards the non-failing state in hypertrophic cardiomyopathy, with cosine shift scores up to ∼2e-2 when candidate therapeutic genes were deleted in-silico. These references further validate the biological relevance of our shift magnitudes and the scISP framework’s ability to benchmark ISP models effectively.

The functional analysis of gene perturbations illustrates how different models recognize various types of gene perturbations. GEARS performs better for ribosomal protein genes and other cellular functions categories and generally demonstrates strengths in multiple categories, once again demonstrating the benefits of supervised training on the Perturb-seq dataset. Geneformer presents balanced detection across categories, while scGPT shows strong precision in ribosomal protein detection but less recognition in other categories. While the reasons for these differences require further investigation and may include technical limitations of CRISPRi technology (e.g. stronger down-regulation of ribosomal genes compared to other genes), this methodology highlights potential differences in the models’ representations. Moreover, these intrinsic differences may provide opportunities to tailor models to specific biological contexts and questions.

The scISP framework is extensible and can be applied to any vector representation of cells per user needs. However, it has limitations. Our study focused on evaluating cell state shifts without analyzing gene embeddings. Gene embeddings can offer insights into gene similarity and potentially build gene interaction networks, providing deeper insights into regulatory mechanisms within cells. Additionally, gene embeddings can help compare the differences in specific genes across different cell states and identify state-specific genes.

Our research focused exclusively on the K562 dataset in CML acquired from large-scale Perturb-seq due to its high quality. Future work should incorporate diverse datasets encompassing various cell types and conditions to further enhance the reliability of the scISP framework and validate the findings presented here. Despite this current limitation, our work represents an initial step towards establishing a unified framework for evaluating ISP models using the cosine shift method.

## Methods

### Ground truth perturbations

The integration of CRISPR-based perturbation technologies with single-cell RNA sequencing (scRNA-seq) has revolutionized genomics research, enabling the generation of large-scale perturbation datasets that capture cellular responses at the transcriptomic level. This approach amplifies the scope of data available per experiment, providing critical resources for training and evaluating deep learning models, as highlighted by Roohani et al.^18^.

To define ground truth perturbations, we applied a rigorous two-tiered approach combining metrics for both on-target interference and transcriptomic impact. On-target knockdown success was assessed by analyzing the differential expression levels of targeted genes, using a t-test to compare perturbed (target) cells with control cells. For efficiency and reproducibility, t-tests were implemented via the rank_genes_groups function within the Scanpy Python package. Perturbations were classified as successful if the targeted gene exhibited a log2 fold change < −1 and an adjusted p-value ≤ 0.05.

To complement the evaluation of on-target effects, we incorporated energy statistics, specifically the E-distance metric introduced by Székely and Rizzo^24^, to quantify transcriptomic alterations across cells. The E-distance captures separation between the overall transcriptomic profiles of target and control cells, offering an unbiased measure of perturbation-induced divergence in gene expression patterns. Building upon previous efforts, such as those by Peidli et al.^19^, this technique identifies perturbations that significantly reshape the cellular transcriptome, enabling a holistic assessment of perturbation success beyond individual gene knockdowns.

Ultimately, strong perturbations—hereafter referred to as ground truth perturbations—were defined as those that achieved both:

1. Successful on-target knockdown (log2 fold change and p-value criteria).
2. Significant transcriptomic disruption, as determined by high E-distance scores.

This dual criterion ensures that the selected perturbations reflect both direct on-target effects and broader cellular transcriptomic disruptions. The resulting ground truth perturbations serve as a robust foundation for evaluating foundational models and machine learning algorithms in their ability to predict and simulate gene perturbation impacts.

### Validation of Cell States

The separation test was designed to determine whether the representations of control and target cell states in the cell representation space sufficiently reflected their biological differences and were statistically distinct. This test evaluated the clustering of control and target states across three models: Geneformer, scGPT, and GEARS. Geneformer and scGPT encode cells as embeddings derived from tokenized gene features, while GEARS represents cells using normalized gene expression values. These models employ different methodologies to represent cell states, and the flexibility of the scISP framework allows validation across a diverse range of representation strategies.

The separation test focused exclusively on ISP-applicable genes, a subset of ground-truth genes for which clustering could reliably be performed. To classify a gene as ISP-applicable, its presence in the input lists for a sufficient number of control cells in the dataset was required (e.g., a minimum of 20 cells), ensuring that it was adequately represented for validation. Geneformer uses input lists of up to 2,048 top-expressed genes to encode each cell, scGPT incorporates up to 1,200 non-zero genes combined with binned values, and GEARS represents cells using 5,000 genes derived directly from transcriptomic data. If the experimentally perturbed gene was absent from the input lists across the required number of control cells in a dataset, it was classified as non-ISP-applicable, disqualifying the gene from inclusion in the separation test and subsequent ISP analysis. These per-cell input limits vary across models, which impacts the number of genes each model can validate in clustering analysis.

To quantify clustering, cosine similarity was chosen as the metric to measure relationships between vector representations due to its reliability and its alignment with the next ISP analysis step, which also relies on cosine similarity to assess perturbation effects. For each perturbation gene, centroids (average vectors) of control and target cell states were calculated within the representation space. Cosine similarity values for control and target cells were measured relative to these centroids: similarities were calculated between control cells and the control centroid (“control-to-control”), control cells and the target centroid (“control-to-target”), target cells and the target centroid (“target-to-target”), and target cells and the control centroid (“target-to-control”). Statistical evaluation relied on one-sided Wilcoxon Rank Sum tests to determine whether control-to-control centroid similarities exceeded control-to-target centroid similarities and whether target-to-target centroid similarities exceeded target-to-control centroid similarities. Genes passing the threshold of adjusted p-values < 0.05 (Benjamini-Hochberg correction) were considered validated, while those failing were excluded from ISP analysis.

This systematic validation approach ensures that scISP can accommodate and fairly compare models with differing approaches to cell representation. Gene representation limits inherently define if a perturbed gene can be included in ISP analysis; thus, validation steps are conducted accordingly to ensure robust benchmarking and meaningful clustering validation.

### ISP Cosine Shift Calculation

Figure 9 illustrates the three key cell states involved in the ISP analysis: control cells, ISP perturbed cells, and in-vitro perturbed target cells. Each dot represents a cell’s embedding or gene expression, modeled differently depending on the approach used by Geneformer, scGPT, or GEARS. The core metric driving the ISP evaluation, the cosine shift, quantifies the movement of ISP perturbed cells toward the target cell state relative to control cells. This calculation ensures that the simulated perturbation effect is directly comparable to experimental in-vitro results.

**Figure 9.**
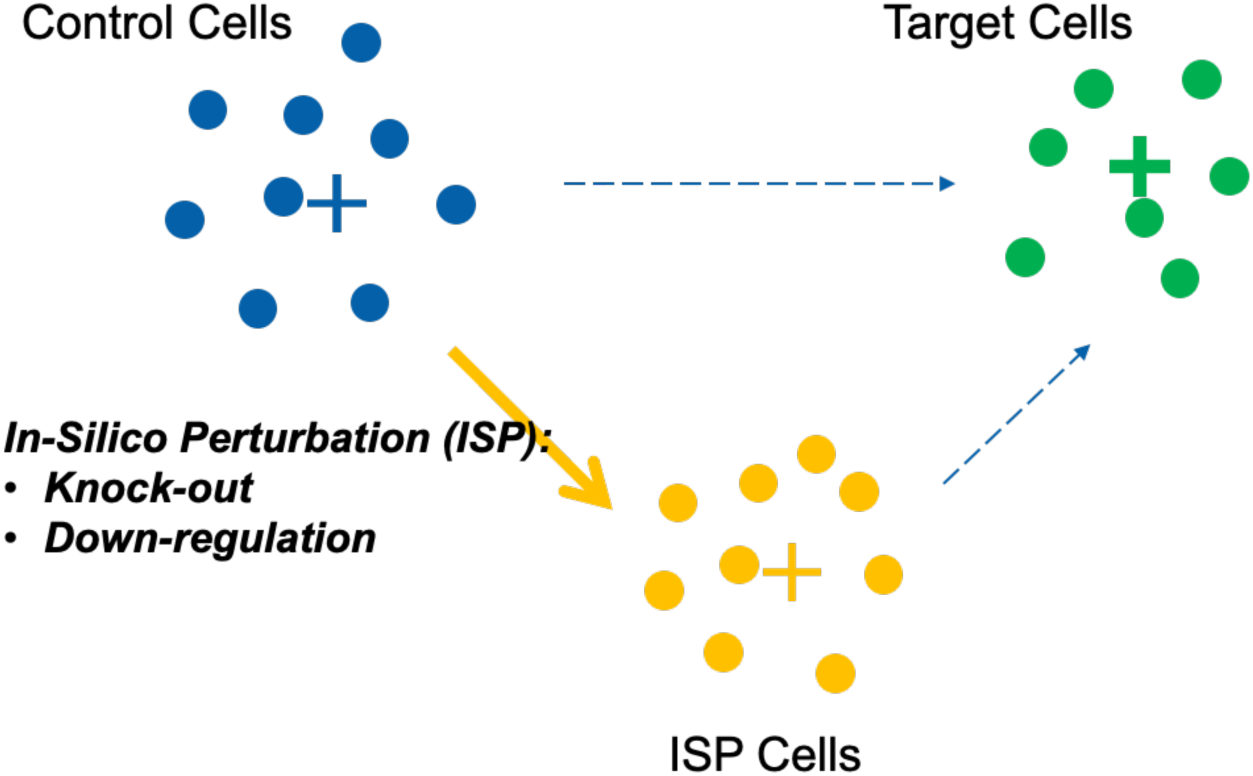
The three cell states of ISP: control cells, ISP cells, and target cells.

The cosine shift for an ISP cell is computed as follows:

#### CS_i_ = cosine(ISP_i_, target_mean) – cosine(control_i_, target_mean)

Here, *CS_i_* represents the cosine shift for the *i*th control cell and its corresponding ISP cell, respectively. The calculation involves the following steps:

- Target Centroid Calculation: For each perturbation gene, we compute the mean embedding of all target cells (*target_mean*) to serve as the reference point for evaluating shifts.
- Cosine Similarity Measurement: The cosine similarity of each ISP cell to the *target_mean* is calculated, followed by the cosine similarity of the corresponding control cell to the same *target_mean*. These values represent each cell’s alignment with the target state.
- Cosine Shift Computation: The difference between the two cosine similarities (ISP cell and control cell) yields a cosine shift value (*CS_i_*) for each cell. Positive shifts indicate movement of ISP perturbed cells toward the target cell state.

### ISP Statistical Testing

To determine the significance of the cosine-shift values, we implemented statistical comparisons between the cosine shifts of ISP cells (sample A) and a baseline set of random shifts (sample B) derived from unrelated perturbation genes. If the shifts in sample A are significantly higher than those in sample B, it demonstrates that the perturbation successfully moved ISP cells closer to the target state.

Given the potential for non-normal distributions of cosine shifts, the Wilcoxon Rank Sum test (Mann-Whitney U test) was selected for comparison, providing a non-parametric approach suitable for broader applicability. Adjusted p-values were calculated using the Benjamini-Hochberg method, ensuring that false discovery rates remain controlled in multiple comparisons.

### Baseline Selection for Rank-Sum Test

To establish a robust baseline, we evaluated several strategies:

- Zero Shifts Baseline: This baseline assumes no perturbation effect, providing a theoretical benchmark for comparison. However, it may not adequately reflect real data variability.
- Down-Sampled Random Shifts Baseline: By down-sampling random shift values to match the size of target cosine shifts, this approach balances comparison scales but introduces variability across iterations, impacting stability and repeatability.
- Combined Random Shifts Baseline: This method uses the entire set of random shifts as the reference, leveraging the larger sample size for reliability and reducing variability.

Ultimately, we selected the combined random shifts baseline for its robustness and stability. However, during analysis, we observed situations where random shifts within certain models, like Geneformer, exhibited a median less than zero. To address this issue, the baseline was adjusted to use the maximum between the random shifts and zero shifts of the same size as the target shifts. This adjustment prevents artificially high pass rates for the rank-sum test, ensuring fair comparison across models.

### Quality Measures

#### Separation Accuracy

Separation Accuracy evaluates whether control and target cell states are distinctly encoded by the model. For each ground-truth gene perturbation, the separation test is performed by assessing within-cluster tightness (how well control cell states cluster together) and between-cluster separation (how far apart control and target cell states are in their representations). These cell representations may exist in embedding spaces for foundation models or transcriptomic spaces for deep learning models like GEARS. Statistically distinct cell states are a prerequisite for meaningful measurement of cosine shift in ISP, as this shift reflects movement from the control state toward the perturbed target state. Overlap or indistinct separation of control and target states would render the concept of “shifting” meaningless. Separation Accuracy is calculated as the percentage of ground-truth gene perturbations successfully passing this test out of the total ground-truth genes, ensuring the model can encode discrete control and target cell states. This measure provides insight into the model’s capacity for preliminary encoding of biologically meaningful shifts between cell states.

#### ISP Accuracy

ISP success was measured by assessing whether perturbed control cells shifted significantly closer to the target cell state based on cosine shifts. For each gene that passed the separation test, cosine shifts were computed, and their significance was evaluated using a rank-sum test with Benjamini-Hochberg (BH)-adjusted p-values. A p-value of ≤0.05 indicated that the perturbation successfully aligned ISP cells closer to the target state, representing a positive prediction. ISP Accuracy was calculated as the proportion of successful predictions out of all ground-truth genes tested.

The denominator for accuracy was set to the total number of ground-truth genes (224) to ensure fairness across models, penalizing those that failed to represent certain genes within their control cell inputs. For instance, Geneformer included 217 of the 224 ground-truth genes in its control cell representations, but its input constraints excluded 7 perturbed genes entirely. Penalizing these omissions reflects limitations in encoding comprehensiveness while avoiding inflated accuracy scores for models that process fewer ISP-applicable genes. This fixed denominator ensures consistent comparisons across all models regardless of differences in input constraints.

#### Mean Reciprocal Rank (MRR)

Additionally, we calculated the Mean Reciprocal Rank (MRR) for each ground-truth gene for each model. For each ground-truth gene *i*, we calculated the cosine shifts of all other genes with respect to the gene’s *target_cell_mean*. This provided a collection of cosine shifts for each gene. We then took the median of this collection. Sorting the genes by their median shifts in descending order, we identified the rank of each gene *i* by locating where its own median cosine shift fell in this sorted list. Iterating through all genes, each gene was assigned a rank. The MRR was then calculated as the average of the reciprocal ranks of these genes. This metric provides a predictive measure of how well models can recommend target genes that were experimentally perturbed, resulting in target cells. Essentially, it evaluates the model’s ability to “guess” which target genes were perturbed based on the observed shifts, akin to a recommendation system. These two measures, ISP Accuracy and MRR, complement each other by providing a comprehensive assessment of model performance in different contexts.

#### Functional categories of gene perturbations

For functional analysis, genes detected by the models were classified into five categories using annotations from Gene Ontology (GO):

1. **Ribosomal Genes**: Genes coding for ribosomal proteins essential for protein synthesis. Ribosomes are essential cellular components that translate mRNA into amino acid sequences. Example genes include RPL24, RPS15A, and RPL34.
2. **Translation Initiation Factors**: Proteins regulating the start of mRNA translation into amino acid sequences. These factors help ribosomes bind to mRNA and ensure the accurate translation start site. Example genes include EIF3H, EIF4E, and EIF2B2.
3. **RNA Binding**: Genes involved in RNA transcription and processing, including regulatory factors for gene expression. Example genes include U2SURP, GTF2H1, and DHX15.
4. **Protein Metabolic Processes**: Genes involved in mitochondrial functions, energy production, and protein regulation pathways. Example genes include IARS2, DLD, and DNAJC19.
5. **Other Cellular Functions**: Genes representing broad or undefined roles across cellular processes. These include signaling, structural components, and protein transport. Example genes include NACA, TSR2, and ABCE1.

Genes overlapping across categories were assigned by prioritizing: Ribosomal Genes > Translation Initiation Factors > RNA Binding > Protein Metabolic Processes > Other Cellular Functions. This framework allowed consistent functional categorization across enrichment analysis, gene interaction mapping, and predictive evaluations.

## Supporting information

Supplemental Enrichment Analysis

## Appendix A Sample UMAP showing Cell States

**Figure A.**
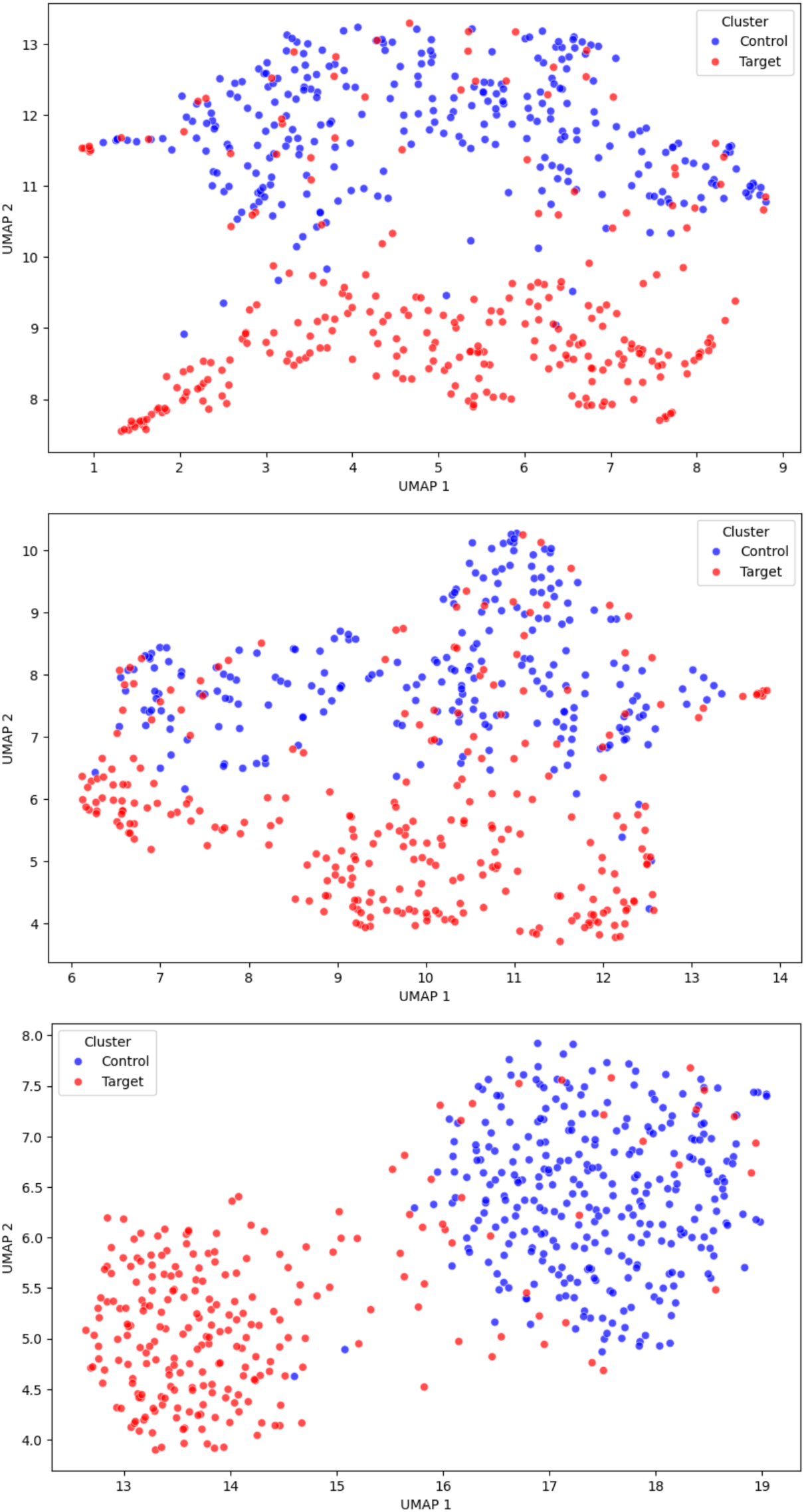
UMAP Visualizations for Gene RPS3. a. Geneformer, b. scGPT, and c. GEARS models illustrating visual separation between control and target cells.

## Appendix B Gene perturbation prediction results

**Table B1.** Geneformer top 20 gene perturbations by cosine shift median.

| Gene | Median cosine shift | Median cosine shift random | Median cosine shift baseline | p_value adj | ISP test result |
| --- | --- | --- | --- | --- | --- |
| RPS3 | 0.000135481 | -3.03984E-06 | 0 | 0 | TRUE |
| RPL14 | 7.37011E-05 | -7.7486E-06 | 0 | 8.6002E-116 | TRUE |
| RPS12 | 6.35684E-05 | -2.563E-06 | 0 | 1.02882E-91 | TRUE |
| HSPA9 | 6.17206E-05 | -2.89083E-06 | 0 | 1.61928E-52 | TRUE |
| RPL9 | 5.78165E-05 | -5.48363E-06 | 0 | 1.2874E-88 | TRUE |
| RPL18 | 5.56707E-05 | -7.09295E-06 | 0 | 9.90999E-75 | TRUE |
| INTS2 | 5.16176E-05 | 4.41074E-06 | 4.41074E-06 | 0.15610711 | FALSE |
| RPSA | 5.03957E-05 | -2.44379E-06 | 0 | 4.0122E-57 | TRUE |
| RPL6 | 3.92497E-05 | -5.72205E-06 | 0 | 1.00415E-38 | TRUE |
| RPL30 | 3.18289E-05 | -8.22544E-06 | 0 | 1.88182E-46 | TRUE |
| HEATR1 | 2.74181E-05 | 2.74181E-06 | 2.74181E-06 | 4.25472E-07 | TRUE |
| RPL11 | 2.66433E-05 | -6.61612E-06 | 0 | 7.49026E-20 | TRUE |
| PTCD1 | 2.58088E-05 | -8.64267E-06 | 0 | 1.24269E-06 | TRUE |
| TSR2 | 2.36034E-05 | -9.53674E-07 | 0 | 6.62955E-19 | TRUE |
| DONSON | 2.32458E-05 | -1.00136E-05 | 0 | 3.31125E-17 | TRUE |
| SFPQ | 2.31564E-05 | 7.15256E-07 | 7.15256E-07 | 2.42845E-07 | TRUE |
| LTV1 | 2.22921E-05 | -1.66893E-06 | 0 | 2.38647E-27 | TRUE |
| EIF3H | 2.16067E-05 | -1.40071E-05 | 0 | 1.31725E-19 | TRUE |
| DHX37 | 2.15769E-05 | -3.8147E-06 | 0 | 6.6373E-20 | TRUE |
| INTS9 | 2.02656E-05 | -8.82149E-06 | 0 | 0.128846368 | FALSE |

**Table B2.** scGPT top 20 gene perturbations by cosine shift median.

| Gene | Median cosine shift | Median cosine shift random | Median cosine shift baseline | p_value adj | ISP test result |
| --- | --- | --- | --- | --- | --- |
| RPL8 | 0.001408011 | 0.000297427 | 0.000297427 | 7.3839E-127 | TRUE |
| RPL36 | 0.001323819 | 0.000353456 | 0.000353456 | 2.68055E-76 | TRUE |
| RPL4 | 0.001305997 | 0.000308692 | 0.000308692 | 1.7969E-101 | TRUE |
| RPL19 | 0.001262844 | 0.000358462 | 0.000358462 | 4.92371E-66 | TRUE |
| RPL9 | 0.001226902 | 0.00030303 | 0.00030303 | 2.14849E-91 | TRUE |
| RPS3 | 0.001087189 | 0.000278056 | 0.000278056 | 8.94812E-64 | TRUE |
| RPL7 | 0.001069069 | 0.000314176 | 0.000314176 | 1.55802E-50 | TRUE |
| RPL23 | 0.001036346 | 0.000396669 | 0.000396669 | 1.87401E-37 | TRUE |
| HSPA9 | 0.001033545 | 0.000151873 | 0.000151873 | 1.97975E-46 | TRUE |
| RPS19 | 0.000971973 | 0.000336707 | 0.000336707 | 6.6827E-37 | TRUE |
| RPL14 | 0.000953019 | 0.000373244 | 0.000373244 | 2.69343E-27 | TRUE |
| RPL6 | 0.000947595 | 0.000390947 | 0.000390947 | 5.07115E-27 | TRUE |
| RPS24 | 0.000904202 | 0.000309587 | 0.000309587 | 1.56796E-29 | TRUE |
| RPS11 | 0.000841081 | 0.000286281 | 0.000286281 | 5.60424E-35 | TRUE |
| RPL18 | 0.000813872 | 0.000323176 | 0.000323176 | 4.70825E-24 | TRUE |
| RPL12 | 0.000786245 | 0.000227451 | 0.000227451 | 1.24196E-25 | TRUE |
| RPL10 | 0.000760764 | 0.000247598 | 0.000247598 | 4.35347E-34 | TRUE |
| RPL37A | 0.000753492 | 0.000289023 | 0.000289023 | 1.58705E-25 | TRUE |
| NACA | 0.000745833 | 0.000375688 | 0.000375688 | 1.50342E-12 | TRUE |
| RPL13 | 0.000730038 | 0.000239134 | 0.000239134 | 2.38614E-28 | TRUE |

**Table B3.** GEARS top 20 gene perturbations by cosine shift median.

| Gene | Median cosine shift | Median cosine shift random | Median cosine shift baseline | p_value adj | ISP test result |
| --- | --- | --- | --- | --- | --- |
| PHB | 0.018918723 | 0.005283773 | 0.005283773 | 9.655E-123 | TRUE |
| RPL31 | 0.016986072 | 0.011063725 | 0.011063725 | 3.36188E-67 | TRUE |
| RPL6 | 0.016982496 | 0.010802537 | 0.010802537 | 3.19096E-64 | TRUE |
| RPL14 | 0.016779244 | 0.01070568 | 0.01070568 | 8.76031E-64 | TRUE |
| RPL34 | 0.016477197 | 0.010326922 | 0.010326922 | 2.62732E-64 | TRUE |
| RPL36 | 0.016105831 | 0.010302216 | 0.010302216 | 2.65703E-70 | TRUE |
| RPL23 | 0.015909523 | 0.01036486 | 0.01036486 | 1.44292E-65 | TRUE |
| RPL18 | 0.015364528 | 0.010095388 | 0.010095388 | 7.3605E-56 | TRUE |
| RPS24 | 0.01518783 | 0.009161592 | 0.009161592 | 4.15432E-60 | TRUE |
| PHB2 | 0.014776856 | 0.004153728 | 0.004153728 | 2.86829E-92 | TRUE |
| RPL7 | 0.01466617 | 0.009325147 | 0.009325147 | 1.63171E-64 | TRUE |
| RPL9 | 0.014422596 | 0.009426802 | 0.009426802 | 4.88468E-57 | TRUE |
| RPS3 | 0.014318198 | 0.008489966 | 0.008489966 | 6.30364E-63 | TRUE |
| RPL4 | 0.014289647 | 0.009524673 | 0.009524673 | 5.92568E-51 | TRUE |
| RPL19 | 0.014173031 | 0.009326756 | 0.009326756 | 6.09491E-62 | TRUE |
| HSPA9 | 0.014006078 | 0.003216654 | 0.003216654 | 1.22831E-96 | TRUE |
| RPL8 | 0.013290852 | 0.00873971 | 0.00873971 | 2.46188E-52 | TRUE |
| RPS7 | 0.012813658 | 0.008162469 | 0.008162469 | 1.13205E-51 | TRUE |
| RPL37A | 0.0128133 | 0.008107275 | 0.008107275 | 6.30938E-57 | TRUE |
| RPS11 | 0.012478113 | 0.007588476 | 0.007588476 | 6.90291E-50 | TRUE |

## Appendix C Annotated Gene List by Functional Category

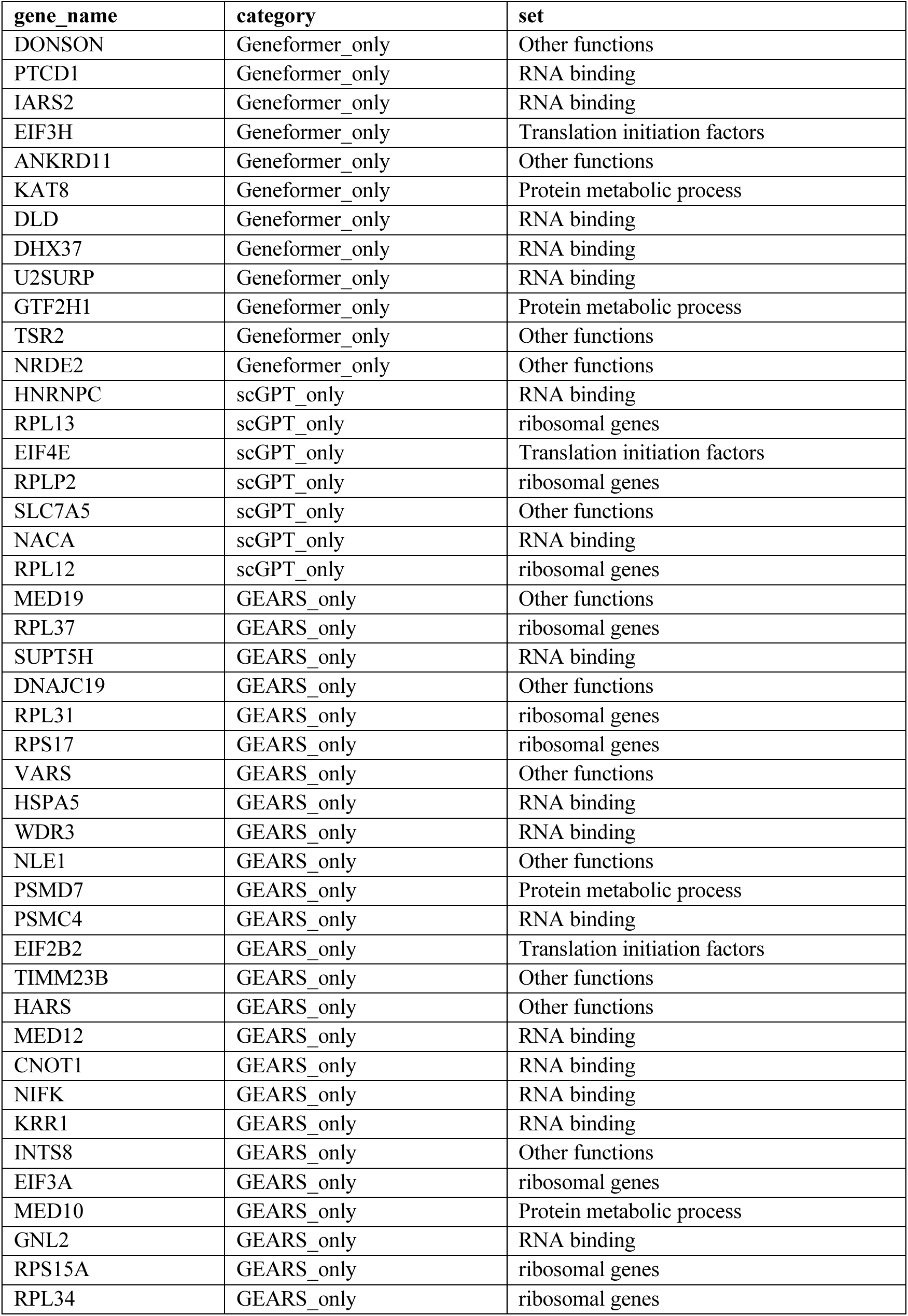
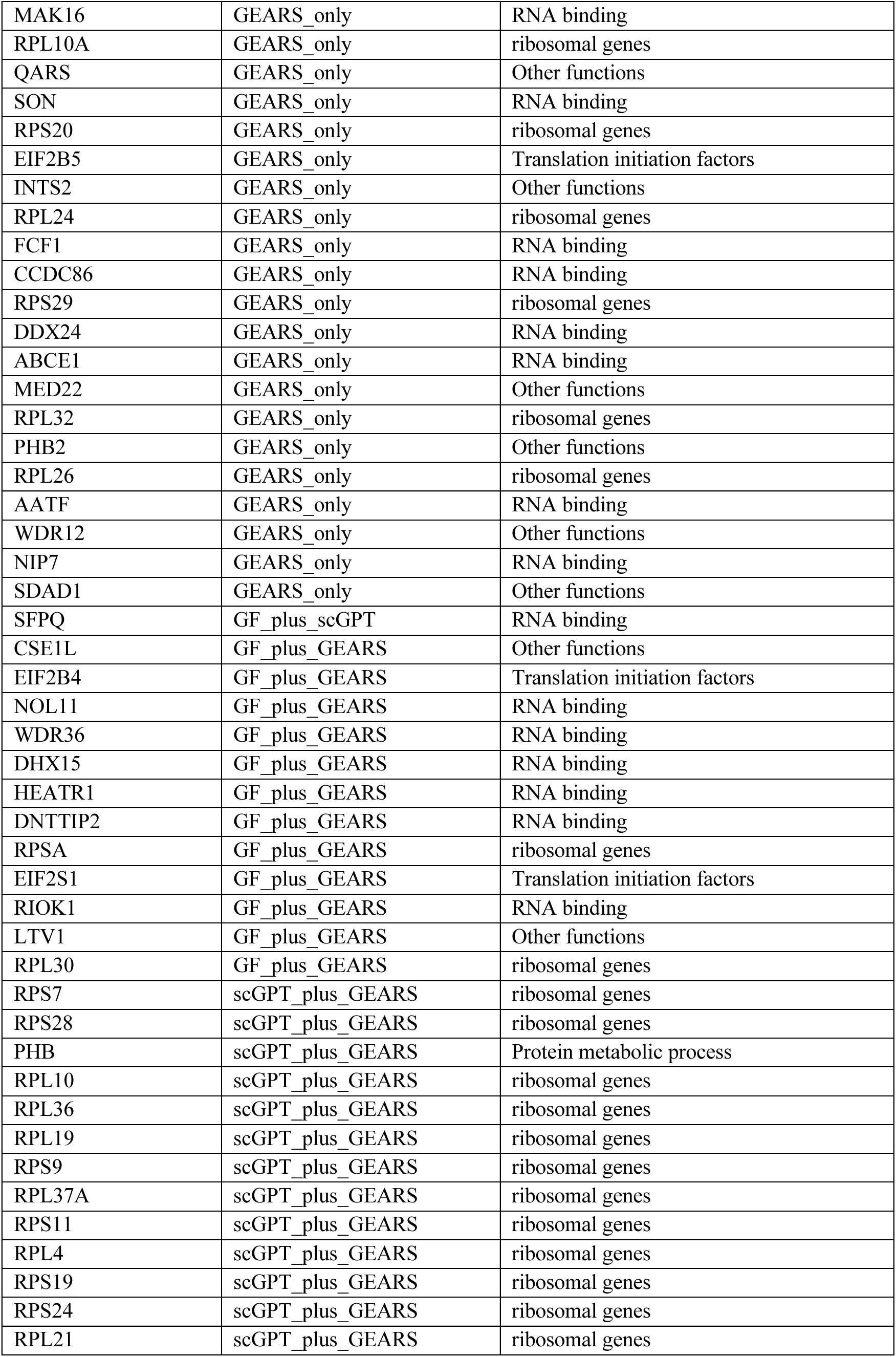
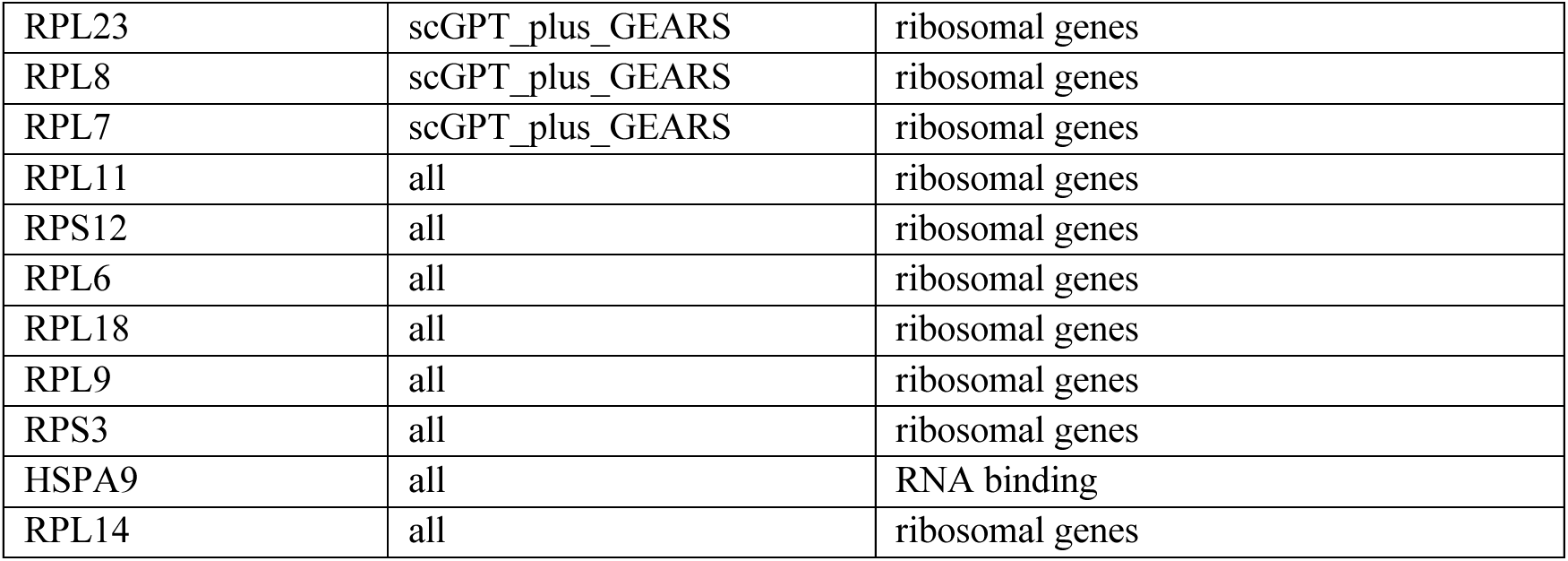

## Appendix D Cosine shifts by functional categories

**Figure D.**
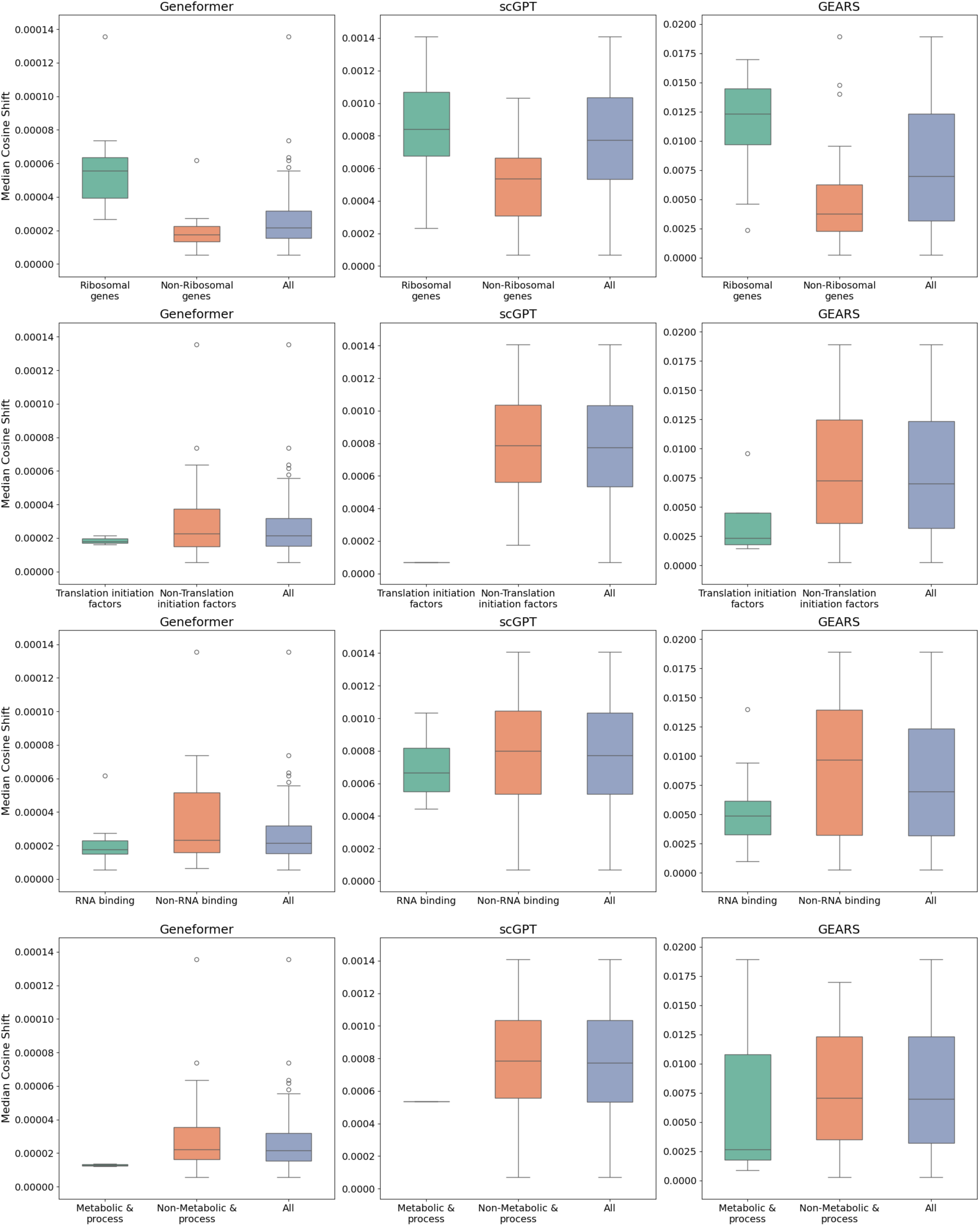
Distribution of Median Cosine Shift Across Gene Categories for Different Models.

